# Role of a versatile peptide motif in controlling Hox nuclear export and autophagy in the *Drosophila* fat body

**DOI:** 10.1101/843383

**Authors:** Marilyne Duffraisse, Rachel Paul, Bruno Hudry, Julie Carnesecchi, Agnes Banretti, Jonathan Reboulet, Leiore Ajuria, Ingrid Lohmann, Samir Merabet

**Affiliations:** IGFL, ENS-Lyon, 32/34 Av. Tony Garnier, 69007 Lyon, France; iVB Parc Valrose, 06108 Nice, France; COS, Im Neuenheimer Feld 230, 69120 Heidelberg, Germany

## Abstract

Hox proteins are major regulators of embryonic development, acting in the nucleus to regulate the expression of their numerous downstream target genes. By analyzing deleted forms of the *Drosophila* Hox protein Ultrabithorax (Ubx), we revealed the presence of an unconventional Nuclear Export Signal (NES) that overlaps with the highly conserved hexapeptide (HX) motif. This short linear motif was originally described as mediating the interaction with the PBC proteins, a generic and crucial class of Hox transcriptional cofactors in development and cancer. Here we show that the HX motif is involved in the interaction with the major CRM1/Embargoed exportin protein. This novel role was found in several *Drosophila* and human Hox proteins. We provide evidence that HX-dependent Hox nuclear export is tightly regulated in the *Drosophila* fat body to control the onset of autophagy. Our results underline the high molecular versatility of a unique short peptide motif for controlling context-dependent activity of Hox proteins at both transcriptional and non-transcriptional levels.

## Introduction

Hox proteins are key developmental regulators that act throughout embryogenesis to specify cell fates and morphogenesis along longitudinal axes in cnidarian and bilaterian animals. These homeodomain (HD)-containing transcription factors (TFs) have specific functions *in vivo* yet recognize highly similar DNA-binding sites *in vitro* (1). It has long been postulated that Hox proteins have to interact with additional transcriptional partners to solve this *in vivo*/*in vitro* paradox (1). The best-characterized cofactors to date are the PBC and Meis proteins, which belong to the TALE-class of HD-containing TFs (2). These cofactors are expressed in many tissues of the embryo and interact with the large majority of Hox proteins (1). They can therefore not account for the full specificity of the Hox regulatory repertoire. In addition, molecular dissections of vertebrate and invertebrate Hox proteins have revealed the important role of the HD and its immediate surrounding region (3), which includes a conserved short linear interaction motif called Hexapeptide (HX). This motif contains an invariant Trp residue that is lying at a variable distance upstream of the HD (4). The Trp residue is important for recruiting the PBC cofactor in dimeric Hox/PBC complexes (1). Hox proteins do however contain a number of additional short peptide motifs that are conserved to different evolutionary extents (3), as well as long disordered regions (5). Recent work showed that human HOX proteins could use various combinations of short peptide motifs, including the HX motif, to interact with the PBC and Meis cofactors, and that long disordered regions could be engaged in non-specific interactions with other types of cofactors (6). These results showed that Hox specific functions can rely on a large and so far poorly investigated portion of their protein sequence. Along the same line, the HX motif was shown to be dispensable for different regulatory functions in Hox proteins (7–9) and functional analyses of the *Drosophila* Hox protein Ultrabithorax (Ubx) bearing a series of small internal deletions highlighted the implication of other protein regions (10).

In order to determine more precisely whether and how other short peptide motif(s) could be involved in the Ubx patterning function, we analyzed a series of Ubx deletions based on structure predictions. This work led to the identification of an unconventional Nuclear Export Signal (NES) that overlaps with the HX motif and that is normally masked by the N-terminal region. The role of this unconventional NES was confirmed during the regulation of autophagy in the larval fat body. The HX motif was also found to be part of a NES in other *Drosophila* and human Hox proteins, highlighting that this divergent function is evolutionary conserved.

Overall our work revealed an astonishing level of functional diversity for a short conserved peptide motif, ranging from transcriptional regulation with generic or tissue-specific cofactors (11) to context-dependent nuclear export. Thus, the HX motif and its immediate surrounding region constitute a privileged platform for diversifying Hox protein function during development and evolution.

## Results and discussion

### N-terminal deletions led to cytoplasmic localization of Ubx

Hox proteins contain a number of short protein motifs and long intrinsically disordered regions that are conserved at different evolutionary extents among the different paralog groups (5, 12). Previous work suggested that the region comprising the HX motif and the HD could serve as a minimal functional Hox module (13, 14), suggesting that the rest of the protein could be dispensable in some instances. Alternatively, work with Ubx underlined that proper specific regulatory activities could also rely on a more distantly located protein motif (10).

To better determine which part of Ubx could be required for early patterning functions in the *Drosophila* embryo, we decided to perform a systematic analysis based on the prediction of the global organization in terms of domains, disorganized regions and short linear motifs by using SLiMPred

(http://bioware.ucd.ie/~compass/biowareweb/Server_pages/slimpred.php). We distinguished six different regions (Fig 1A) that could be analyzed individually or in combination upon N- and C-terminal deletions (Fig 1A). The resulting deleted forms were N-terminally fused to the C-terminal fragment of the fluorescent Venus protein (VC; EV1) for subsequent protein interaction analyses (see below) and functionally tested in the epidermis with the UAS/Gal4 system (15). As expected, ectopic expression of the wild type VC-Ubx fusion protein generated A1-like segments in the thorax (Fig 1B). The small C-terminal deletion (VC-Ubx^dC^) did not notably affect the ability of Ubx to induce A1-like denticle belts in the thorax (Fig 1B). By comparison, removing the first 235 residues of Ubx, (VC-Ubx^dN235^ construct) completely abolished the A1 specification function (Fig 1B). The same result was obtained when the dN235 deletion was combined with the dC deletion (VC-Ubx^dN235dC^ construct; Fig 1B), suggesting that the C-terminal portion is quite neutral in this context. Therefore, the two other smaller N-terminal deletions were also analyzed in the absence of the C-terminal region (EV1). Results showed that the corresponding constructs (VC-Ubx^dN183dC^ and VC-Ubx^dN130dC^) were also inactive, as well as the HD only (construct VC-Ubx^dN282dC^, Fig 1B). Together, these results confirm that the long N-terminal portion is required for A1-patterning function of Ubx.

**Figure 1.**
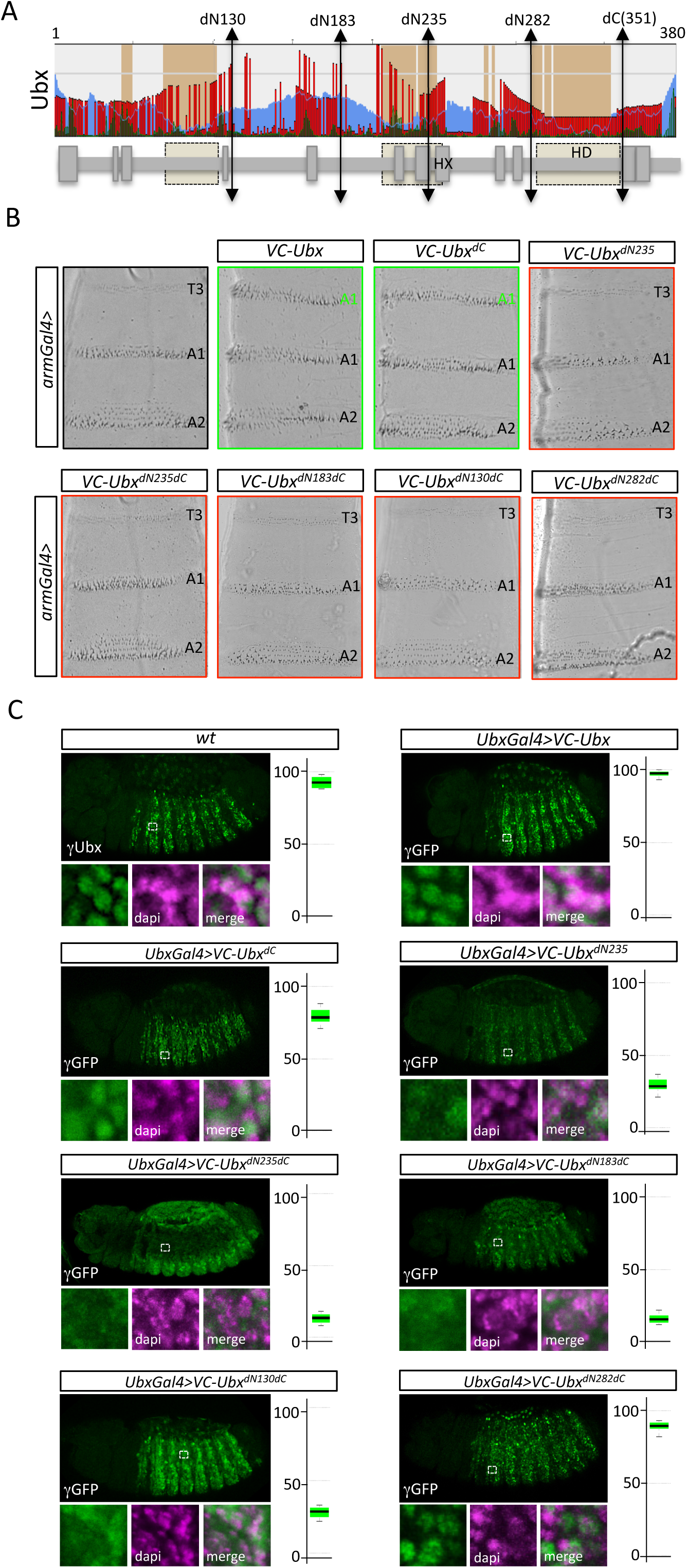
N-terminal deletions lead to cytoplasmic localization of Ubx in the embryo. **A.** Schematic representation of Ubx (adapted from SLiMPred). Prediction of short linear interaction motifs (SLiMs, green peaks), disordered regions (blue waves), ordered domains (brown boxes) and level of conservation of each residue (red peaks) is shown along the protein sequence. N- and C-terminal deletions are indicated, as well as the Hexapeptide (HX) motif and the Homeodomain (HD). A simplified representation of Ubx is shown below the global structure. Boxes denote SLiMs. **B.** Cuticles of larvae expressing the different Ubx constructs with the *armadillo(arm)-Gal4* driver, as indicated. Ubx constructs are fused to the C-terminal fragment of the fluorescent Venus protein (VC) for subsequent analyses. Acquisitions are centered on the thoracic T3 and abdominal A1 and A2 segments. Ubx is responsible of the specification of A1, which is characterized by few rows of refringent denticle belts. T3 denticle belts are much less refringent and specified by Antennapedia (Antp), while A2 has more refringent denticle belts with a global trapezoidal shape organization (under the specification of Abdominal-A). Ectopic expression of wild type or C-terminal form of Ubx induces the transformation of T3 into A1, as attested by the denticle belts. Ectopic expression of N-terminal deleted forms of Ubx has no effect on T3 denticle belts, indicating that these constructs are not able to promote Ubx-like transformation. **C.** Immunostaining of endogenous Ubx and VC-Ubx constructs (green), as indicated. VC-Ubx constructs were expressed with an *Ubx-Gal4* driver and revealed with an anti-GFP antibody recognizing the VC epitope, as previously described (28). Immunostainings were co-revealed with dapi (magenta) to stain nuclei. Enlargements are shown below each illustrative confocal acquisition of the embryo. Quantification of Ubx or GFP staining is provided on the right as a percentage of nucleus/cytoplasmic signals. The N- and to a lesser extent C-terminal deletions induced a cytoplasmic localization of VC-Ubx constructs.

We then performed immunostaining to verify that each construct was correctly expressed in the nucleus. Quantification was performed with induced expression levels close to the endogenous Ubx expression level in the A1 segment, as previously done (16). Surprisingly, we found that deletions could impact on the correct nuclear localization. Full-length Ubx construct was expressed like endogenous Ubx, with an exclusive nuclear localization (Fig 1C). In contrast, the C-terminal deleted form was slightly less localized in the nucleus (around 78%) and found also present in the cytoplasm (Fig 1C). The dN235 deletion led to a stronger decrease of nuclear localization, with only 30% of the construct in the nucleus on average (Fig 1C). Combining the dN235 and dC deletions further decreased the nuclear fraction (around 20%: Fig 1C), with a corresponding increase of the cytoplasmic fraction. The two other deleted constructs (VC-Ubx^dN183dC^ and VC-Ubx^dN130dC^) were also mostly cytoplasmic (Fig 1C). Finally, the minimal HD-containing form of Ubx (VC-Ubx^dN282dC^) was strongly localized in the nucleus (Fig 1C). This last result suggests that nuclear localization signal(s) (NLS) could be present within the HD of Ubx, in accordance with previous observations (17). In contrast, the cytoplasmic localization of N-terminally deleted forms could be due to the presence of Nuclear Export Signals (NES) that are inactive in the context of full length Ubx. More particularly, the respective cytoplasmic or nuclear localization of VC-Ubx^dN235dC^ and VC-Ubx^dN282dC^ constructs indicates that the region comprised in between the two N-terminal deletions could be instructive for the cytoplasmic localization.

### Identification of an atypical Nuclear Export Signal (NES) that overlaps with the HX motif in Ubx

To identify putative NES(s), we scanned the Ubx protein sequence with two different algorithms, LocNES (http://prodata.swmed.edu/LocNES/LocNES.php) and NetNES (http://www.cbs.dtu.dk/services/NetNES/). Surprisingly, NetNES did not predict any NES with a significant confidence score (EV2). LocNES identified three putative NES with high or low confidence scores; two of which are in the N-terminal region (residues 37-50 and 191-205) and the other one in the HD (residues 318-332; EV2). Importantly, neither NetNEs nor LocNES predicted a putative NES in the region discriminating exported (Ubx^dN235dC^) and non-exported (Ubx^dN282dC^) Ubx constructs.

Although NESs are of variable nature, they all contain 4-5 hydrophobic residues (often Ile/Leu/Met/Phe/Val) arranged in a particular pattern (18). NESs can still adopt diverse conformations and have been classified in ten different categories (18). Interestingly, we found that the Ubx protein region comprised in between the dN235 and dN282 deletions contained a sequence that could resemble to a NES but in an inverted orientation (<u>L</u>DSRIKGAIA<u>M</u>: Fig 2A). Inverted NES sequences have been described in some instances and classified as class1a-R NESs (18, 19). Moreover, this putative NES is conserved in flies (EV3), with the last hydrophobic residue lying in the core HX motif (Fig 2A).

**Figure 2.**
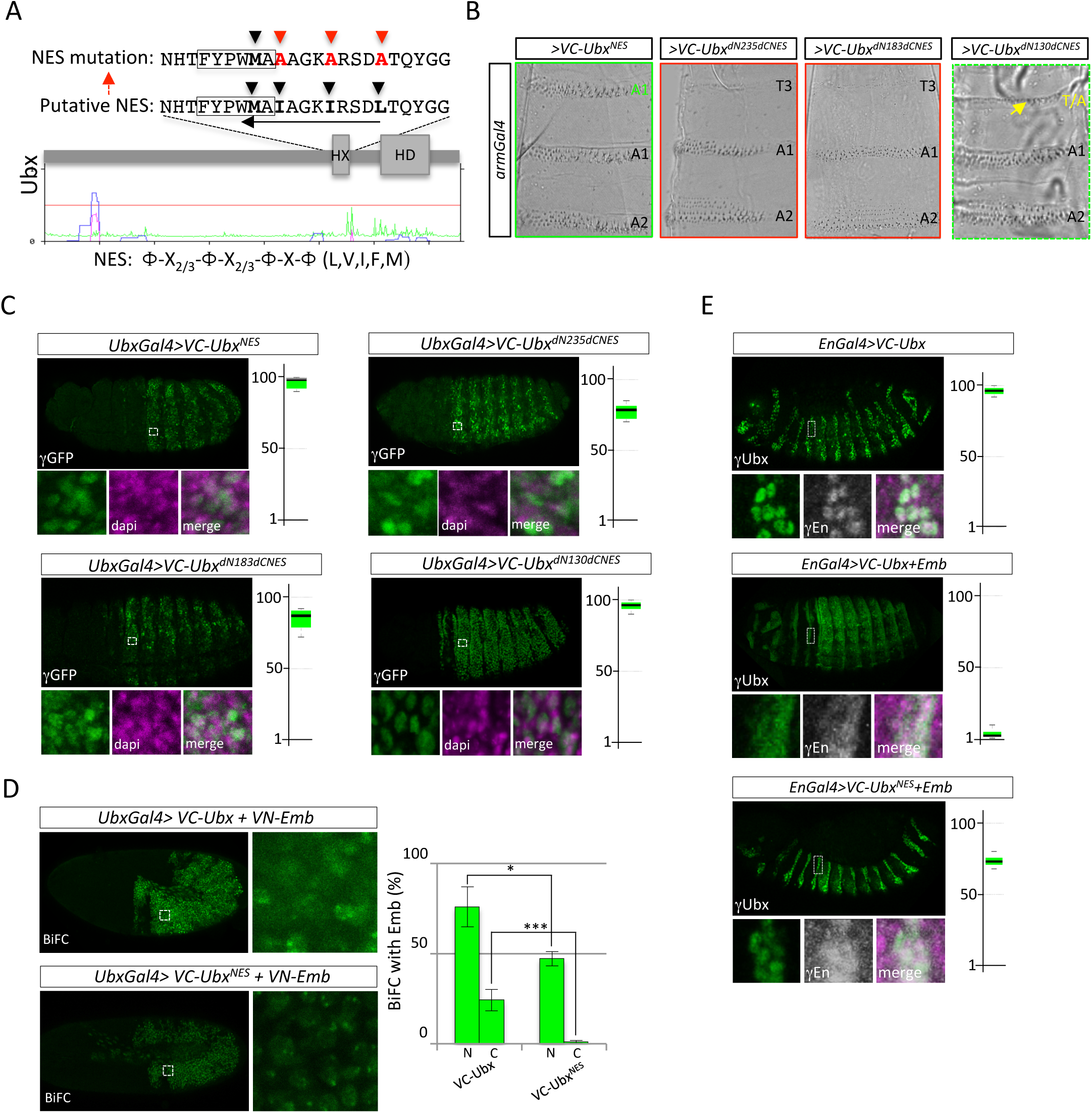
Identification of an unconventional Nuclear Export Signal (NES) in the HX-containing region of Ubx. **A.** Sequence of the HX-containing region in Ubx. The HX motif is boxed and hydrophobic residues that could participate to the NES are indicated (black arrowheads). The consensus NES is given below the predicted graph for full Ubx (from NetNES). No putative NES is predicted along the Ubx sequence. Mutations generated to abolish the putative NES are indicated (NES^m^: red arrowheads). These mutations do not touch the HX motif residues. Black arrow shows the orientation of the putative NES. **B.** Cuticles of larvae expressing the NES-mutated forms of Ubx, as indicated. The NES mutation does not affect the ability of Ubx to induce ectopic A1-like segments. The NES mutation does also not rescue the loss of activity of the deleted forms. Only the dN130dC-deleted form shows weak activity, with a mixed thoracic-abdominal-like transformation in T3 (yellow arrow depicting the few refringent abdominal-like denticles). **C.** Quantification of NES mutated forms in the nucleus, as previously described in Fig 1. The NES mutated forms are all located back in the nucleus. **D.** Bimolecular Fluorescence Complementation (BiFC) between wild type or NES-mutated Ubx and Embargoed (Emb). Fusion proteins are fused to the N-(VN) or C-(VC) terminal fragment of Venus and expressed with *Ubx-Gal4*, as indicated. BiFC was quantified in the nucleus (N) and cytoplasm (C) of stage 10 live embryos. Graph on the right show the statistical quantification. The NES mutation affects the interaction with Emb. **E.** Forced co-expression of Emb induces NES-dependent cytoplasmic localization of Ubx. The VC-Ubx fusion proteins (green) were expressed in posterior parasegments with the *engrailed (en)-Gal4* driver, with or without cold HA-tagged Emb, as indicated. Quantification of VC-Ubx in the nucleus was performed with an anti-Ubx antibody in the T3 segment (absence of endogenous Ubx). The effect of Emb is also visible with anti-En staining (gray), which becomes cytoplasmic in the presence of Emb. Mutation of the NES renders Ubx insensitive to the effect of Emb.

To assess whether this sequence works as a NES in Ubx, we mutated the first three hydrophobic residues into Ala residues, as classically done for NES mutations (20). These mutations were intentionally chosen to avoid disturbing the integrity of the HX motif. The effects of these mutations were analyzed in the context of full length and deleted forms of Ubx. Functional assays showed that the simple mutation of this putative NES did not alter the ability of Ubx to induce ectopic A1-like segments (VC-Ubx^NES^, Fig 2B). The NES-mutated forms of VC-Ubx^dN183dC^ and VC-Ubx^dN235dC^ constructs were still inactive (Fig 2B), while the NES-mutated VC-Ubx^dN130dC^ construct induced an intermediate phenotype with the formation of weak A1-like denticles in the thorax (Fig 2B). Immunostaining revealed that all the mutated constructs were strongly expressed in the nucleus, with a strong decrease (constructs VC-Ubx^dN235dCNES^ and VC-Ubx^dN183dCNES^) or complete absence (construct VC-Ubx^dN130dCNES^) of localization in the cytoplasm (Fig 2C). These observations thus confirmed that the inverted NES that we identified was responsible for the cytoplasmic localization of the deleted forms of Ubx. In addition, it also demonstrated the long N-terminal region is absolutely required for the A1 specification function of Ubx.

Nuclear export is ensured in *Drosophila* by the major exportin called Embargoed (Emb, (21)), or CRM1 in vertebrates (22–25). To assess whether the newly identified NES motif in Ubx could be important for recruiting Emb, we performed a series of different complementary interaction assays. First, we performed Bimolecular Fluorescence Complementation (BiFC, (26)) between full length Ubx fused to the C-terminal fragment of Venus (construct VC-Ubx) and Emb fused to the complementary N-terminal fragment of Venus (construct VN-Emb). BiFC was performed with wild type or NES-mutated Ubx. Fusion constructs were expressed with the *Ubx-Gal4* driver and BiFC was analyzed in the live embryo, as previously described (27, 28). Results showed that Ubx could interact with Emb, both in the nucleus and the cytoplasm (Fig 2D). In comparison, BiFC with NES-mutated Ubx was strongly affected; with a complete loss of fluorescence in the cytoplasm and 40% decrease in fluorescent intensity in the nucleus (Fig 2D).

The specific and NES-dependent activity of Emb was confirmed independently of BiFC, by co-expressing VC-Ubx with cold HA-tagged Emb: in this context, the forced co-expression of Emb was sufficient to induce the cytoplasmic localization of Ubx (Fig 2E). Importantly, this cytoplasmic localization was not observed when the NES-mutated form of Ubx was co-expressed with Emb (Fig 2E), indicating that the Emb-dependent export of Ubx was fully dependent on the integrity of the newly identified NES.

Finally, physical interaction between Ubx and Emb was also assessed by GST-pull down experiments by using *in vitro* produced GST-Ubx, wild-type or NES-mutated, and S2R+ cells extracts expressing Emb (see Materials and Methods). Control experiments showed that the NES mutation did not affect the interaction with the Exd cofactor (as expected) while no interaction could be observed with the GFP (EV4). Results showed that Ubx and Emb could directly interact and that the pull-down of Emb associated with Ubx was significantly reduced when the NES was mutated (around 30% loss: EV4).

### The HX motif is acting as a NES in other Drosophila and non Drosophila Hox proteins

Given its conservation among the Hox family members, we wondered whether the HX motif could be part of a putative NES in other *Drosophila* Hox proteins. As previously done with Ubx, we used available scripts based on published NES sequences to scan several *Drosophila* (Deformed, Dfd; Sex combs reduced, Scr; Antennapedia, Antp; AbdominalA, AbdA; AbdominalB, AbdB) and human (HOXB4, HOXA5, HOXA7 and HOXA9) Hox proteins. This analysis did not reveal any consensus NES sequence with high confidence in the HX-containing region, except for AbdB, which has a derived HX motif (corresponding to a single W residue; EV5-6). Surprisingly, our own analysis again revealed the presence of a putative NES in the HX-containing region of Dfd and Scr (Fig. 3A and 3D; EV7). In contrast to Ubx, these putative NES sequences are not inverted, belonging to the class-1a, and they fully overlap with the HX motif. Moreover, the hydrophobic residues of these putative NESs are widely conserved among invertebrate and vertebrate species, suggesting they could be of functional significance.

**Figure 3.**
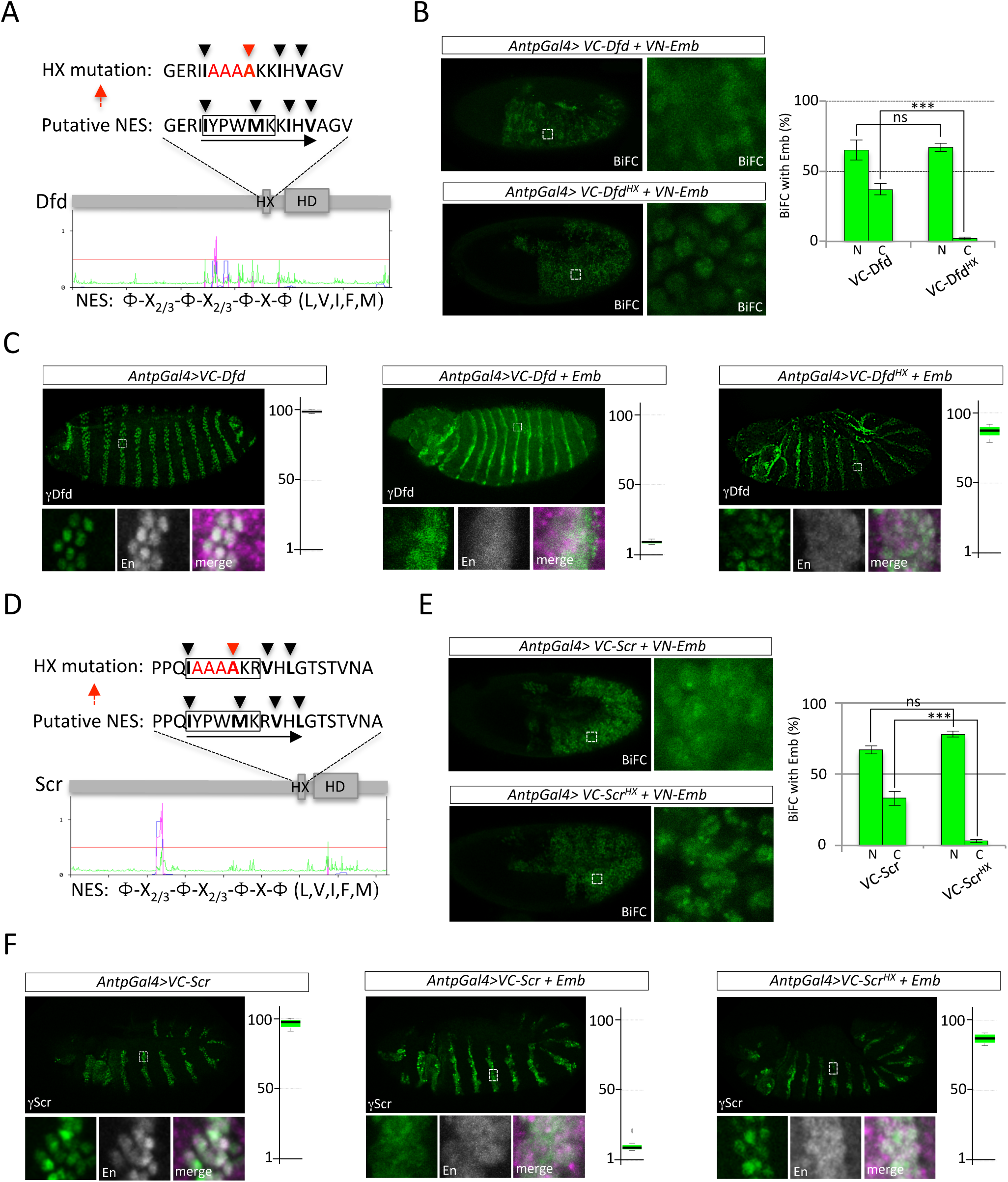
The HX motif is part of an unconventional NES in other Hox proteins. **A.** Protein sequence of the HX-surrounding region in Deformed (Dfd). Black arrowheads indicate hydrophobic residues that could participate to the NES. Black arrow shows the orientation of the putative NES. The classical HX mutation (highlighted in red) affects the core of this unconventional NES. The graph below illustrates predicted NES with NetNES, taking into account the consensus repartition of hydrophobic residues (indicated below the graph). NetNES predicts a putative NES with a good confidence score in the region upstream of the HX (magenta peak). **B.** BiFC between wild type or HX-mutated VN-Dfd fusion protein and VC-Emb, as indicated. Fusion proteins are expressed with the *Antennapedia (Antp)-Gal4* driver. Graphs on the right show the statistical quantification of BiFC signal in the nucleus (N) and cytoplasm (C) of epidermal cells of stage 10 live embryos. The HX mutation affects the Dfd-Emb interaction only in the cytoplasm. **C.** Genetic interaction between Dfd and Emb. The co-expression of cold HA-Emb triggers Dfd in the cytoplasm except if the HX motif is mutated. Quantification is performed as described in the Figure 2. **D.** Protein sequence of the HX-surrounding region in Sex combs reduced (Scr). Black arrowheads indicate hydrophobic residues that could participate to the NES. Black arrow shows the orientation of the putative NES. The classical HX mutation (highlighted in red) affects the core of this unconventional NES. The graph below illustrates predicted NES with NetNES, taking into account the consensus distribution of hydrophobic residues (indicated below the graph). NetNES predicts a putative NES with a good confidence score in the region upstream of the HX (magenta peak). **E.** BiFC between wild type or HX-mutated VC-Scr fusion protein and VN-Emb, as indicated. Fusion proteins are expressed with the *Antennapedia (Antp)-Gal4* driver. Graphs on the right show the statistical quantification of BiFC signal in the nucleus (N) and cytoplasm (C) of epidermal cells of stage 10 live embryos. The HX mutation affects the Scr-Emb interaction only in the cytoplasm. **F.** Genetic interaction between Scr and Emb. The co-expression of cold HA-Emb triggers Scr in the cytoplasm except if the HX motif is mutated. Quantification is performed as described in the Figure 2.

The putative role of the NES of Dfd and Scr was assessed as previously done with Ubx. We first performed BiFC with Emb, using wild type or HX-mutated versions of each Hox protein (16). Results showed that the HX mutation, which affected only one hydrophobic residue of the putative NES (Fig 3A and 3D), abolished BiFC in the cytoplasm but had no effect on the nuclear Hox-Emb interaction (Fig 3B and 3E). Co-expression of wild type or HX-mutated Hox proteins with cold Emb also confirmed that the cytoplasmic localization induced by Emb was fully dependent on the HX integrity (Fig 3C and 3F). These results thus confirm that the HX motif of Dfd and Scr is required for Emb-dependent cytoplasmic localization.

Similarly, we observed that the cytoplasmic localization of human HOXA5 induced by the co-expression of human CRM1 in HEK-293T cells was also fully dependent on the integrity of the HX motif (EV8). Together, these results demonstrate that the HX motif is part of a functional NES in some but not all Hox proteins, highlighting that the NES function of the HX-containing region was acquired independently in different Hox proteins and animal species.

To further validate that the HX motif could work as an autonomous NES, we fused the corresponding sequence of Ubx or Scr upstream of the Dronpa fluorescent protein (EV9). Control constructs with full length Ubx or a consensus NES sequence (29) fused to Dronpa confirmed the localization of the fluorescent reporter in the nucleus or the cytoplasm, respectively (EV9). Importantly, the fusion of Dronpa with the HX motif of Ubx or Scr led to its cytoplasmic localization with the same efficiently as consensus NES sequence (EV9), validating that these peptide sequences could act as autonomous NES.

### Active nuclear export of Ubx is necessary for inducing autophagy in the larval fat body

Hox proteins are strongly expressed in the nucleus of embryonic cells, suggesting that the nuclear export is actively inhibited during embryogenesis. Interestingly, previous work noticed that Hox proteins were absent in the fat body cells of L3-wandering (L3-W) larvae, while they were present in fat body nuclei at the earlier feeding stages to repress the expression of autophagy-related genes (*atg*) and block autophagy process (30). We confirmed that Ubx was present in the nucleus of fat body cells until late L3-feeding (L3-F), and that no or very low nuclear Ubx proteins could be detected at the L3-W stage (Fig. 4A-A’). The absence of nuclear Ubx (and other Hox proteins) coincides with the onset of developmental autophagy, highlighting the importance of Hox nuclear clearance for the temporal control of autophagy. Although it was described that Hox genes were transcriptionally repressed (30), we hypothesize that Hox nuclear clearance could also rely on active nuclear export at the L3-W stage. To test this hypothesis, we used a clonal system to abolish the expression of *emb* by RNAi only in few cells in an otherwise wild type tissue, as previously described (30). Results showed a nuclear retention of endogenous Ubx at the L3-W stage, exclusively in the *emb* mutant cells, but not in surrounding wild type cells (Fig. 4B-B’). We also observed retention of the Atg8-mCherry reporter in the nucleus, with the absence of cytoplasmic autophagosomes, suggesting that the Atg8-mCherry reporter was not fully repressed but could not be properly exported in the absence of Emb (Fig. 4B-B’).

**Figure 4.**
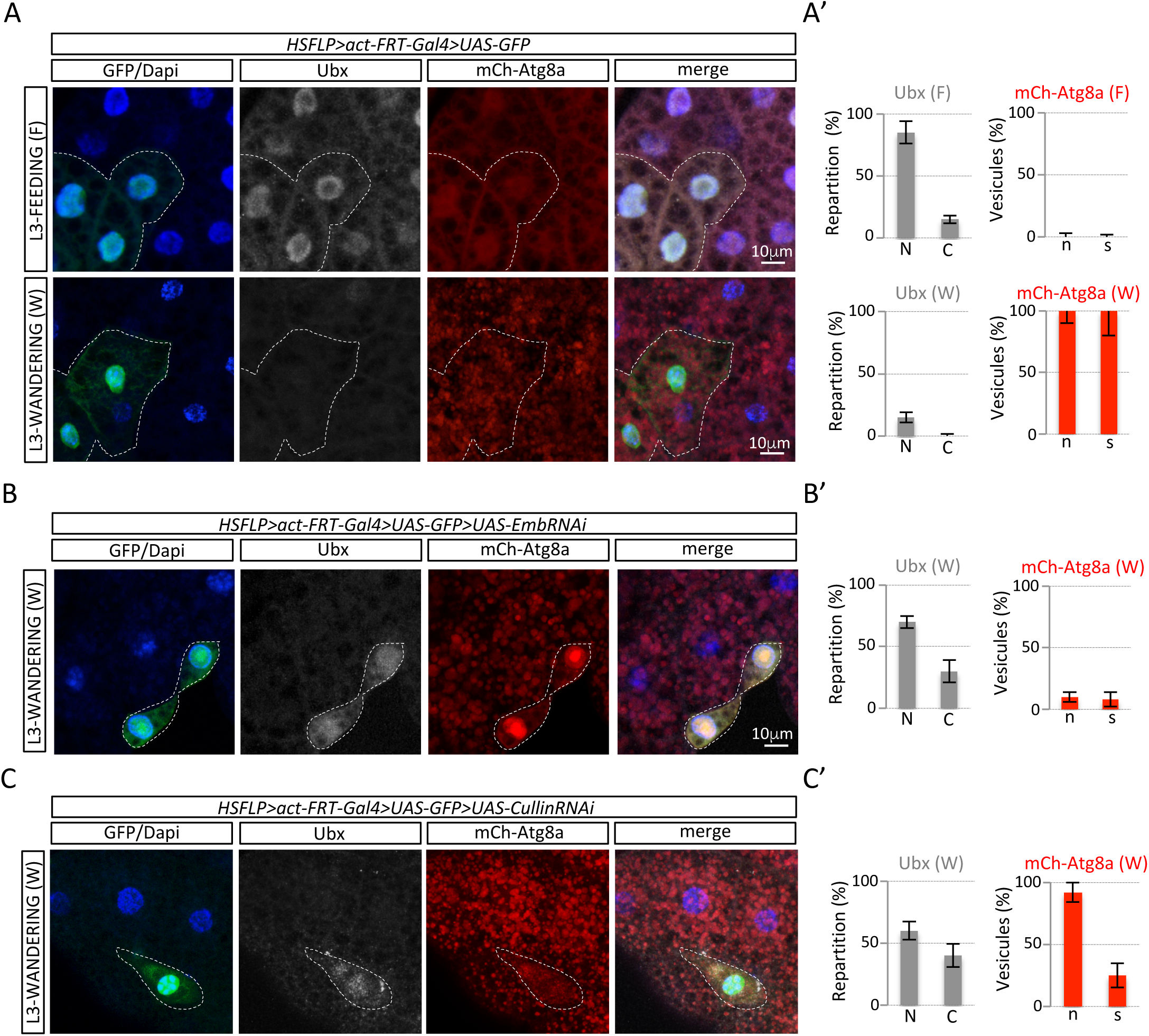
Ubx is actively exported from fat body nuclei and degraded in the L3-wandering (W) stage. **A.** Immunostaining of endogenous Ubx (gray) in the fat body of L3-Feeding (F) and L3-W larva. Cloned are recognized with the GFP expression (green). The ATG8-mCherry reporter (red) allows following the off (weak nuclear expression) and on (dotted staining corresponds to autophagosomes) states of autophagy in the fat body of L3-F and L3-W larvae, respectively. **A’.** Graphs showing the statistical distribution of Ubx staining in the nucleus (N) or cytoplasm (C), together with the number (n) and size (s) of ATG8-mCherry vesicles in fat body cells (see also Materials and Methods). **B.** Blocking the nuclear export by inhibiting the expression of Emb with RNAi leads to nuclear accumulation of Ubx in the nucleus of L3-W fat body cells. Note that ATG8-mCherry also accumulates in the nucleus. **B’.** Statistical quantification of Ubx and Atg8-mCherry distribution as described in A’. **C.** Blocking the proteasome activity by the expression of *Cullin* RNAi reveals accumulation of Ubx in L3-W fat body cells. **C’.** Statistical quantification of Ubx and Atg8-mCherry distribution as described in A’.

The absence of Ubx in the cytoplasm of L3-W fat body cells suggests that the protein is degraded once it is exported from the nucleus (Fig. 4A). To verify that Ubx was indeed exported into the cytoplasm at L3-W stage, we inhibited the protein degradation machinery. This was achieved through the inhibition of the Cullin (Cul) ubiquitin ligase expression: clonal cells expressing RNAi against *Cul* showed endogenous Ubx expression at the L3-W stage, which coincided with a significant reduction of the size of Atg8-mCherry autophagosomes (Fig. 4C-C’). This result reveals that Ubx nuclear export leads to the cytoplasmic degradation of the protein. Stabilizing Ubx in the cytoplasm could allow the protein to return into the nucleus, therefore repressing back the expression of *atg* genes that eventually affected the formation of autophagosomes.

Altogether these observations highlight that active Emb-dependent nuclear export is occurring in the transition from L3-F to L3-W fat body tissue to exclude Hox proteins from the nucleus and release the transcriptional repression of *atg* genes.

### The atypical NES and the N-terminal region control nuclear export of Ubx in the larval fat body

Analyses in the epidermis of the early embryo showed that the activity of the inverted NES found in Ubx could only be revealed in the absence of the N-terminal region. The same constructs were analyzed in the L3-W fat body to assess whether the nuclear export could rely on the same intra-molecular regulatory loops. In parallel, the effect on the nuclear export was functionally correlated at the level of developmental autophagy regulation by looking at the ATG8-mCherry reporter expression. Constructs were expressed using the same clonal system as previously described to allow side-by-side comparison of the effect with the surrounding wild type cells.

As expected (30), the wild type VC-Ubx construct was strongly expressed in the nucleus and led to significant repression of autophagy (Fig. 5A-A’). Interestingly, mutating the NES led to even stronger expression in the nucleus and increased repression of autophagy (Fig. 5A-A’). This observation suggests that the inverted NES of Ubx might have a physiological role in the fat body. Deleting the C-terminal part induced a slight reduction of the nuclear fraction and a corresponding slight decrease in autophagy repression when compared to full length Ubx (Fig. 5A-A’). The additional deletion of the first 235 residues (construct Ubx^dN235dC^) was sufficient to strongly affect the nuclear localization (70% loss on average) and to abolish the effect of Ubx on repressing autophagy (Fig. 5A-A’). This result shows that the N-terminal region, and to a lesser extent the C-terminal region, inhibit the activity of the NES in the fat body. Accordingly, mutating the NES in the context of the dN235 and dC deletions (construct VC-Ubx^dN235dCNES^) was sufficient to retain the protein in the nucleus and actively repress autophagy (Fig. 5A-A’). In contrast, the minimal form containing the HD (construct VC-Ubx^dN282dC^) was not able to repress autophagy although it was fully expressed in the nucleus (Fig. 5A-A’).

**Figure 5.**
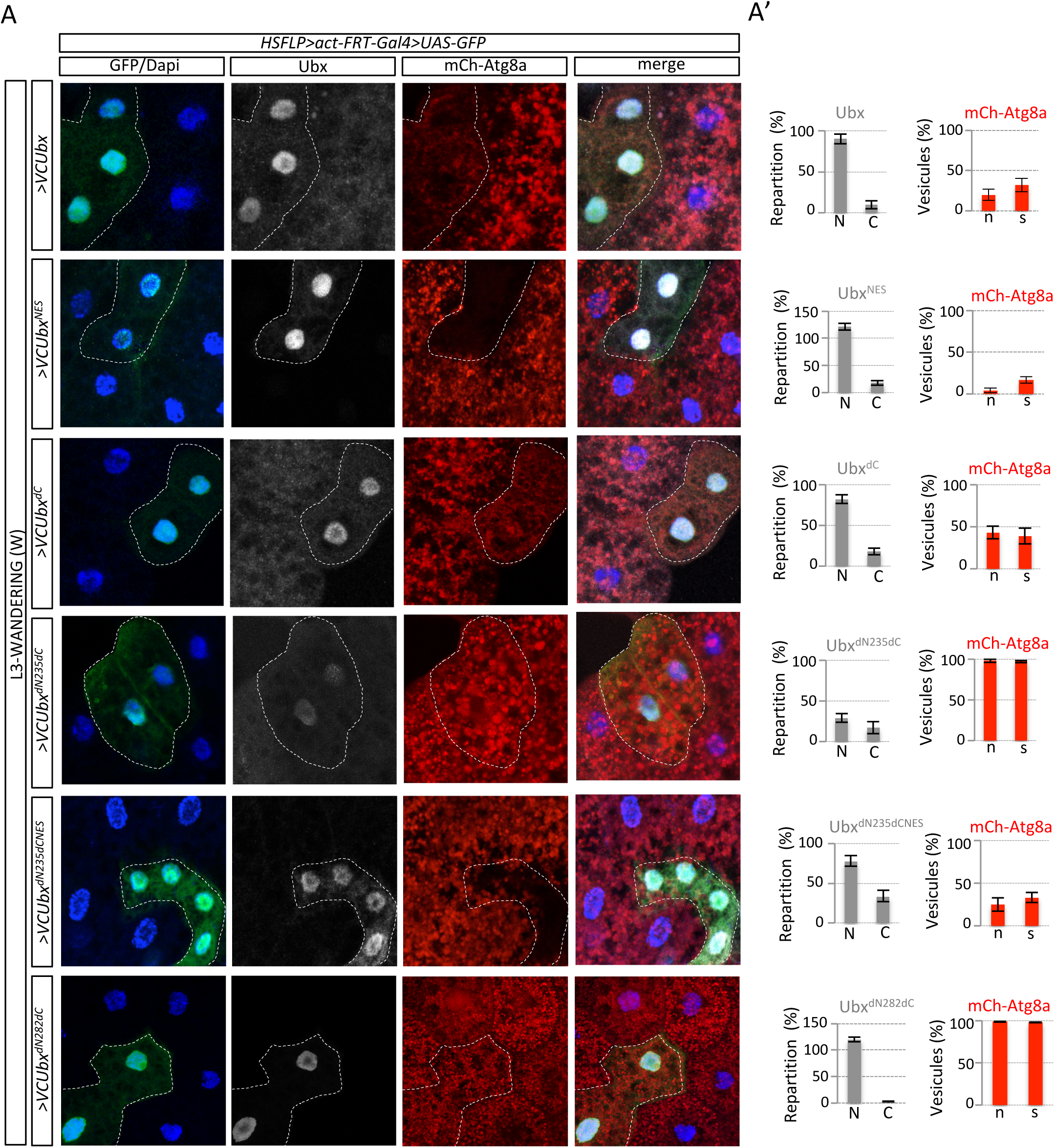
The non-canonical NES controls Ubx nuclear export in the *Drosophila* fat body. **A.** Cloned are recognized by the GFP expression (green) and Ubx constructs are revealed with an anti-Ubx (full length Ubx constructs) or anti-HA (truncated Ubx constructs) antibody (gray). Dapi (blue) stains for nuclei and ATG8-mCherry for autophagy (red). **A’.** Graphs showing statistical distribution of VC-Ubx constructs and ATG8-mCherry, as described in the Figure 5. The NES mutation blocks the nuclear export induced by the dN235dC deletion, allowing the constructs to reside in the nucleus and inhibit the autophagy. The minimal dN282dC form is localized in the nucleus but is not able to inhibit autophagy.

Altogether, these observations confirmed that the inverted NES identified from analyses in the embryo was also used and negatively controlled by the N-terminal region. Combining N-terminal deletions with the NES mutation showed that the molecular determinants necessary for Ubx activity were however not identical in the different developmental contexts. Deleting the first 130 residues of Ubx affected its ability to promote the formation of A1 denticle belts in the embryo, while deleting the first 235 residues did not affect the ability of Ubx to repress autophagy in the larval fat body. These observations underline that Ubx functions are relying on different molecular modules and that previously described synthetic Hox peptides (13, 14) could likely ensure only part of the versatile functions of Hox proteins *in vivo*.

The N-terminal part of Ubx contains long intrinsically disordered regions (IDRs), which are inherent molecular features of Hox proteins (5). These regions are more generally described to be under alternative splicing and post-translational modifications, allowing SLiMs to engage context-specific interactions with surrounding partners (31). Here, an important role of the disordered region resides in its ability to mask the NES, keeping the Hox protein active for engaging interactions in the nucleus. This observation underlines that IDRs could positively influence the interaction potential of a regulatory protein by regulating its subcellular location and in particular in the case of Ubx by inhibiting the recruitment of CRM1/Emb. Several post-translational modifications have been described to either promote or inhibit the activity of NES in TFs. Among them are the phosphorylation (32), sumoylation (33), O-GlcNac modification (34) and acetylation (35, 36). These post-translational modifications could potentially be involved in the NES masking activity of the N-terminal region. Along this line, Ubx is phosphorylated throughout embryogenesis (37) and HOX proteins are described to interact with the CBP/p300 acetyl transferase (38). Such post-translational modifications could potentially be involved in the NES-masking activity of the Ubx N-terminal region.

Finally, the finding that the HX motif could be part of a NES was surprising since this motif is working as a docking platform for the PBC-class cofactors in the context of Hox/PBC dimeric complexes. This role is however dispensable in the presence of the third Meis partner, which is required for the nuclear translocation of PBC proteins (39). The dispensability of the HX motif in the context of Hox/PBC/Meis trimeric complexes was observed for the majority of Hox proteins studied so far (6), suggesting that the HX motif could be available for other types of interactions. Accordingly, the HX motif was described to promote or reversely inhibit the interaction with different types of TFs, and this dual activity was tissue-specific (40). The role of the HX motif as part of a docking platform for the Emb/CRM1 exportin protein enlarges the repertoire of its putative binding partners. All together, these observations highlight the astonishing level of context-dependent protein interaction flexibility for this short motif (EV10). These inherent molecular properties may explain that the HX motif has been conserved in the large majority of Hox proteins in animals, underlining that sequence conservation is not only compatible but also indispensable for functional diversity. Such molecular diversity implies a tight control from the other surrounding protein regions, explaining that the same short peptide motif could have distinct functions in different protein contexts (41).

## Figure legends

**Expanded View 1. Schematic representation of the deleted Ubx constructs under study.** Symbols are as in Figure 1. The VC fragment is shown (green).

**Expanded View 2. Prediction of NES in Ubx. A.** Protein sequence of Ubx. The HX motif (red box), HD (blue box) and UbdA (UA, back box) motif are indicated. **B.** Sequences of predicted NES with LocNES. The corresponding putative NES are underlined and in italic in the full Ubx sequence. **C.** No NES is predicted with high confidence (only magenta peaks above the red line are considered as significant) when using NetNES.

**Expanded View 3.** Evolutionary tree of Diptera, showing that the non-canonical NES (highlighted in green) is only conserved in Ubx proteins of fruit flies (Tephritidae (T) and Drosophilidae (D) branches) from the Brachycera sub-order. This NES is not present in Ubx proteins from the Culicidae (C) family (mosquitos).

**Expanded View 4. Interaction NES-dependent between Ubx and Emb *in vitro*. A.** Immunoblots of GST-pull-down assay using *in vitro* produced GST-fused derivatives (GST alone; GST-Ubx-wild type, WT; GST-Ubx-NES-mutated, NES) and whole S2R+ cell extracts expressing VC-Emb (A), GFP (B) or Flag-Exd (C). GFP antibody was used to recognize VC-Emb and GFP. Flag antibody was used to recognize Flag-Exd. Inputs (Ip 2%) are systematically loaded, as indicated (first lane of each gel). Pull-down assays showed that Ubx-WT and Ubx-NES could interact with Emb and Exd but not with GFP (lanes 3-4). Quantification of interactions relative to GST-Ubx signal indicated a decrease of 30% of the interaction of Emb with Ubx-dNES when compared to the Ubx-WT form.

**Expanded View 5. Analysis of *Drosophila* and human Hox protein sequences with NetNES.** The sequence of putative NESs is provided when the prediction score was good enough (magenta peak above the red line threshold). No NES was predicted in Antp and AbdB. NES were predicted in the N-terminal region of Dfd and Scr, and in the homeodomain (HD) of AbdA, HOXB4, HOXA5, HOXA7 and HOXA9.

**Expanded View 6. Analysis of *Drosophila* and human Hox protein sequences with LocNES.** The sequence of putative NESs is provided when the prediction score was good enough. NESs were predicted only in Antp, AbdA and AbdB. Of note, one of the NES predicted in AbdB contains the derived HX motif (corresponding to a single W residue: bolded and red).

**Expanded View 7. Conservation of the non-canonical NES covering the HX motif in Dfd (A) and Scr (B) proteins from invertebrate and vertebrate species.** *Dm: Drosophila melanogaster. Am: Apis melifera. Bm: Bombyx mori. Tc: Tribolium castaneaum. Csa: Cupiennus salei. Mm: Mus musculus. Dr: Danio rerio. Nv: Nasonia vitripennis.* Red arrowheads highlight hydrophobic residues of the putative NESs.

**Expanded View 8. The HX motif of human HOXA5 is part of a NES mediating CRM1-dependent export in live HEK293T cells. A.** Sequence of the putative non-canonical NES overlapping with the HX motif of HOXA5. The HX mutation is indicated (red). Prediction of NES with NetNES is shown below the schematized protein (with one canonical NES in the HD). **B.** The wild type and HX-mutated fusion HOXA5-mCherry proteins are localized in the nucleus upon transfection in HEK cells. Co-expression of CRM1 induced cytoplasmic localization of HOXA5-mCherry but not of HX-mutated HOXA5-mCherry. This effect was observed in 100% of co-transfected cells. Cells expressing CRM1 are recognized by the GFP that is fused downstream of CRM1 (see Materials and Methods).

**Expanded View 9. HX-containing peptides from Ubx and Scr behave as autonomous NES when fused to the fluorescent Dronpa protein. A.** Schematic representation of the different constructs, with the peptide sequence fused to Dronpa in each case. Fusion with a consensus NES or full length Ubx was used as a control. **B.** Expression of fusion fluorescent proteins in live *Drosophila* embryos. Illustrative confocal capture is provided in each case. Enlargements are shown on the right. Fusions are expressed with the *Ubx-Gal4* driver, together with NLS-mCherry to label the nuclei. The Ubx-Dronpa fusion protein is expressed in the nucleus, as expected. In contrast, the fusion to a consensus NES leads to the cytoplasmic localization of Dronpa. Peptides containing non-canonical NES from Ubx and Scr also induce cytoplasmic localization of Dronpa, as efficiently as the consensus NES peptide.

**Expanded View 10. The HX motif is involved in different types of molecular interactions to control context-dependent activity of Hox proteins.** The HX motif was originally described for its generic role in recruiting the PBC cofactor in the context of Hox/PBC complexes (left panel). The HX motif was also described to promote or inhibit interactions between Hox proteins and different transcription factors in the *Drosophila* embryo. This function is tissue-specific, enabling the Hox protein to establish specific interaction networks in different cell contexts (central panels: (40)). This work described a novel tissue- and stage-specific (L3-W fat body)) function of the HX motif in participating to the Hox nuclear export through the interaction with the exportin CRM1/Emb (right panel).

## Materials and Methods

### Fly stocks

The different *Gal4* drivers used are: *armadillo(arm)-Gal4* (BL1560), *Ubx-Gal4*^*M1*^ (42), *engrailed(en)-Gal4* (43), *Antp-Gal4* (16). *UAS-RNAi* constructs were from Vienna (*UAS-EmbRNAi, P(KK102552)*) or Bloomington (*UAS-Cul1RNAi, P(Trip.HM05197)*) stock centers.

Different UAS-driven fly lines were previously generated: *UAS-VC-Ubx, UAS-VC-Scr, UAS-VC-Scr*^*HX*^ (16). The following UAS-driven fly lines were generated in this study: NES- and/or deleted forms of *UAS-VC-Ubx, UAS-HA/VC/VN-Emb, UAS-VN-Dfd*/*Dfd*^*HX*^, *UAS-Dronpa* constructs. All constructs were sequence-verified before transgenesis.

### Flip-out expression of UAS constructs and mitotic clones and quantitation of protein localization and autophagy

Clonal expression and quantifications in fat body cells were performed as previously described (30). Briefly, males carrying UAS constructs were crossed to *yw,hs-Flp;r4-mCherry::Atg8a;Act>CD2>GAL4,UAS-GFPnls* females. Crosses were kept for 1 night for egg laying at room temperature. Thereafter, parents were removed and the tube was incubated at 25°C for 3 days. Leaky expression of the heat-shock inducible Flp led to random ‘‘flip out’’ of the CD2 cassette, allowing Gal4 to be expressed and to activate the expression of UAS-driven transgenes and nuclear GFP (to identify the cellular clones). mCherry-Atg8a (mCh::Atg8a) was used for tracing autophagy activity in the whole fat body.

Quantification of protein localization in the nucleus and cytoplasm and autophagy were performed with ImageJ. Dapi was used to stain for nuclei and to differentially quantify the fluorescence between the nucleus and the cytoplasm. Threshold was adjusted to select dots for the red channel, and identical areas were chosen randomly in clones and neighboring wild type control cells. Selected areas were analyzed (analyze particles, show, masks), and the counts and average sizes were noted from the summary of masked images. At least five images from 5 animals and from three independent experiments were quantified for each genotype. The quantification number and size data were summarized in Excel and normalized to control (average fluorescence, dot number and size of control cells were set to 100%).

### Preparation of Samples and Imaging of Fixed Tissues

Larvae were dissected 4 days (L3 Feeding stage) or 6 days (L3 Wandering stage) after egg laying. Larval fat bodies were fixed at room temperature for 20 min in PBS, 3.7% formaldehyde (formaldehyde methanol free, Thermo scientific) and washed for 30 min in 1xPBS. All incubations were done in PBS, 2% BSA, 0,1% Triton X-100 at 4°C following standard procedures.

Images were acquired with a Zeiss LSM780 confocal microscope. Pictures were edited with ImageJ and Adobe Photoshop CS6 Version.

### Cell culture and transfection

S2R+ Drosophila cells were maintained at 25°C in Schneider medium supplemented with 10% FCS, 10U/ml penicillin and 10µg/ml streptomycin. Cells were simultaneously seeded and transfected with Effectene (Qiagen) according to the manufacturer’s protocol. For interaction assay, 10.10^6^ cells were seeded in 100mm dishes. Cells were harvested in Phosphate Buffered Saline (PBS) after 48 hours of transfection and pellets were resuspended with NP40 buffer (20mM Tris pH7.5, 150mM NaCl, 2mM EDTA, 1% NP40) supplemented with protease inhibitor cocktail (Sigma-Aldrich) and 1mM of DTT and treated with Benzonase (Sigma).

Transfections in HEK cells were performed by using the JetPRIME reagent (Polyplus), with a total amount of 2 mg of DNA: 1 mg of the HOXA5-mcherry and 1 mg of empty or CRM1-GFP-containing PcDNA3 vector. Coverslips were taken 20h after transfection, which allows having fluorescence level below saturation with each condition. Analysis was performed with a Zeiss LSM780 confocal microscope. Pictures correspond to the Z projection of stacks, using the Zen software. Four to six different fields of cells were acquired under the same confocal parameters at the 20x objective from two independent experiments in each condition.

### Constructs

All UAS constructs were sequence verified before fly transformation and transgenic fly lines were established with the ΦC31 integrase (44). All UAS-Hox constructs were inserted on the same landing site on the second chromosome (22A3), as the previous published UAS-Hox constructs (16). UAS-Emb constructs were inserted on the third chromosome (62E1). HOXA5-mCherry, HOXA5^HX^-mCherry and CRM1-GFP were cloned in the PcDNA3 vector and are under the *CMV* promoter for expression.

### Cuticle preparation and Immunohistochemistry

Embryo collections, cuticle preparations and immunodetections were performed according to standard procedures.

The following primary antibodies used were: chicken anti-chicken anti-GFP (ab13970, Abcam, 1/500), rabbit anti-GFP (A11122, Molecular Probes, 1/500), mouse anti-Ubx (FP3.38, DSHB 1/100), rat anti-HA (11867423001, Roche, 1/500).

Fluorescent secondary antibodies used were: Alexa Fluor 488 (A11039; A11008, Molecular Probes), Cy-3 (A1051, Molecular probes), Alexa Fluor 647 (A21235, Invitrogen). Vectashield with DAPI (Vector Labs) was used to stain nuclear DNA.

### SDS-Page and Immunoblotting

For western blot analysis, proteins were resolved on 8 to 15% SDS-PAGE, blotted onto PVDF membrane (Biorad) and probed with specific antibodies after saturation. The antibodies (and their dilution) used in this study were: GFP (Life Technologies, 1/3000), GST (Cell signalling, 1/5000e), Flag-M2 (Sigma, 1/1000e).

### Protein purification and GST-Pull down

Wild type and NES-mutated GST-tagged Ubx proteins were cloned in pGEX-6P plasmids and sequence verified before using. GST-tagged Ubx proteins were produced from BL-21 (RIPL) bacterial strain, purified on Gluthatione-Sepharose beads (GE-healthcare) and quantified by Coomassie staining. *In vitro* interaction assays were performed with equal amounts of GST or GST fusion proteins in affinity buffer (20mM HEPES, 10μM ZnCl2, 0.1% Triton, 2mM EDTA) supplemented with NaCl, 1mM of DTT, 0.1mM PMSF and protease inhibitor cocktail (Sigma). 300µg of S2R+ whole cell lysates were subjected to interaction assays for 2h at 4°C under mild rotation. Bound proteins were washed 4 times and resuspended in Laemmli buffer for western-blot analysis. Input fraction was loaded as previously described (45).

### BiFC visualisation in live embryos

BiFC analysis was performed as previously described (27). Briefly, over night embryos were kept at 4°C for 24 h before live imaging. Living embryos were dechorionated and mounted in the halocarbon oil 10S (commercialized by VWR, Pennsylvania, USA). Images of BiFC in stage 10 live embryos were acquired with a Zeiss LSM780 confocal microscope, using adjusted Venus excitation wave length (500nm) and emission filters (535nm). Identical parameters of acquisition were applied between the different genotypes. The number and intensity of the all pixels (for each embryo) were measured using the histogram function of the ImageJ Software.

## Aknowledgements

We thank the Bloomington and Vienna stock centers for providing the *Drosophila* lines, and the Developmental Studies Hybridoma Bank for antibodies. Work in S. M.’s laboratory was supported by the CNRS, ENS Lyon, Fondation pour la Recherche Médicale and Centre Franco-Indien pour la Promotion de la Recherche Avancée.

